# Learning interpretable compositional features associated with continuous covariates

**DOI:** 10.64898/2026.11.04.764349

**Authors:** Xingyi Chen, Stephanie C. Hicks

## Abstract

Compositional variation as a function of continuous covariates, such as time, presents challenges for learning interpretable features and accounting for their construction in subsequent inference. Motivated by temporal change in isoform usage in genomics, we introduce IsoVita, a statistical framework integrating covariate-informed balance learning designed for compositional data with complementary tests for abrupt and gradual compositional variation. IsoVita estimates a dominant covariate-dependent direction in the log-ratio coordinates and approximates it as a log ratio of their geometric mean proportions between two learned component groups, linking statistical evidence to explicit component membership. Change point and spline tests evaluate this interpretable feature, while a selection-aware permutation procedure repeats feature construction and testing under each permutation. In transcript-level simulations, IsoVita maintained low empirical false discovery proportions with power to detect differences and change point accuracy improving as temporal signals strengthened. However, the performance depended on trajectory shape and study design. Applications to two human brain bulk RNA-sequencing datasets identified widespread temporal variation in isoform composition while retaining transcript-level interpretability. IsoVita integrates interpretable compositional feature learning with inference along continuous covariates, with potential applications beyond transcriptomics. IsoVita is available as a PyPI package (https://pypi.org/project/isovita).

## 1 Introduction

Continuous covariates often reveal patterns that comparisons between predefined groups may obscure, including abrupt transitions, gradual trends, or non-monotone trajectories. Such patterns arise over time, across exposure levels and environmental gradients, and along inferred pseudotime. Methods such as Slingshot and tradeSeq reconstruct trajectories and test association variation, primarily at the gene level for single-cell RNA-sequencing [1, 2]. Compositional measurements, such as microbiome data, pose an additional challenge: their components convey relative information and must be interpreted jointly [3, 4]. However, there lack methods to identify compositional variation associated continuous covariates.

Transcript isoform usage is another an example of compositional data and relative transcript abundances can reveal regulatory variation beyond gene expression [5]. Methods for differential transcript usage and alternative splicing include DEXSeq, DRIMSeq, LeafCutter, and SPIT [6–9]. Methods including IsoDOT, CompDTU, satuRn, Gaussian-process models, and Iso-maSigPro [10–14] also accommodate continuous covariate or temporal structure. Recent work with IsoGraph summarizes within-gene isoform usage in log-ratio coordinates and integrates usage and gene abundance into networks of coordinated variation [15]. A complementary goal is to learn an explicit contrast between component groups along a continuous covariate and account for that learning when testing association.

Log-ratio, or balances, methods provide a foundation for analyzing relative variation, with applications ranging from geological compositions to microbial communities [3, 16]. Balances contrast geometric means of two component groups, making their interpretation explicit through component membership [4, 17]. Principal balances support dimension reduction [17], while supervised methods such as selbal, coda4microbiome, and LCoDaCoRe learn interpretable signatures for prediction from cross-sectional or longitudinal data [16, 18, 19]. We instead seek a balance summarizing variation along a continuous covariate with tests sensitive to abrupt and smooth changes. Because the covariate guides both feature construction and testing, inference must account for feature selection [20].

Here, we introduce IsoVita, a framework combining covariate-informed balance learning with selection-aware inference. IsoVita approximates a dominant fitted direction in log-ratio coordinates using a contrast between two learned component groups, allowing other components to remain unused. Change point and spline tests identify abrupt and smooth variation, respectively. A permutation procedure repeats feature construction and testing to account for their dependence [21, 22] with multiple-testing adjustment applied separately to each test family [23]. Transcript-level simulations evaluate feature condensation, empirical false discoveries, statistical power, change point localization, and sensitivity to trajectory shape and sampling design. Applications to dorsolateral prefrontal cortex and hippocampus RNA-sequencing data examine temporal variation in isoform usage, building on previous studies investigating transcriptional signatures in the human brain [24–27]. Together, these analyses illustrate a general approach to learn interpretable compositional features and to test their variation along continuous covariates.

## 2 Results

### 2.1 Interpretable compositional features associated with continuous covariates

IsoVita learns an interpretable compositional feature and tests its variation along a continuous covariate. We use temporal isoform trajectories to motivate the problem: predefined groups obscures within-group variation, transition timing, and trajectory shape, event when they preserve between-group differences (**Figure 1**). Feature condensation instead summarizes coordinated changes while retaining the continuous axis (**Figure 1B**). IsoVita learns a contrast between component groups that serves a common feature for complementary association tests.

**Figure 1:**
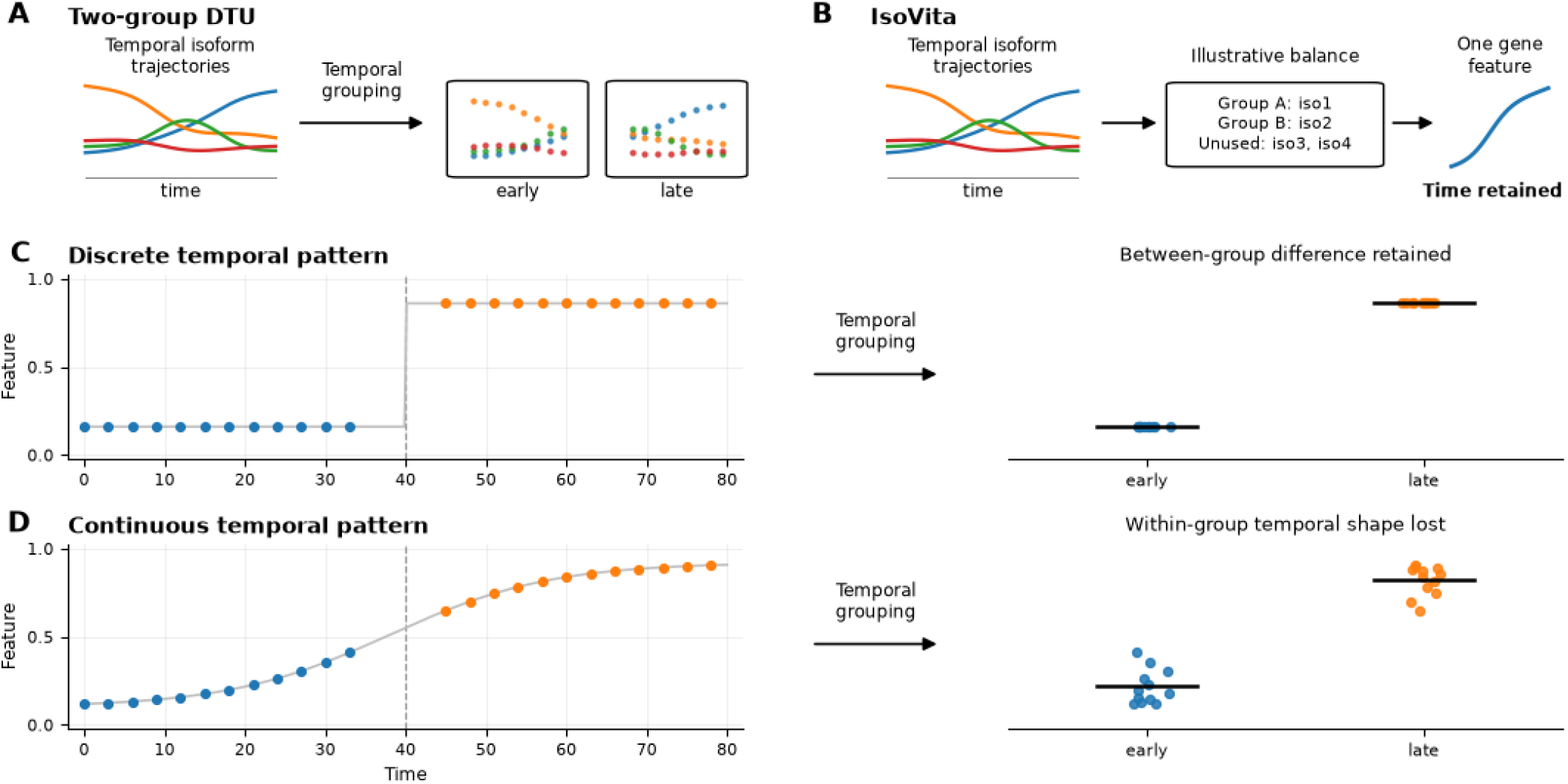
Interpretable compositional features associated with continuous covariates. (**A**) Conventional two-group differential transcript usage (DTU) analysis replaces a continuous time variable with predefined early and late groups. (**B**) IsoVita condenses the same isoform trajectories into one interpretable gene-level log-ratio feature while retaining time information. The component groups shown illustrate balance construction. (**C**) For a discrete trajectory with a step at time 40, grouping retains a between-group difference but removes information about the transition’s location. (**D**) For a continuous trajectory, grouping retains a between-group difference but obscures the within-group temporal shape. Colors in A and B distinguish between isoforms. Blue and orange points in C and D represent early and late samples, respectively. Black horizontal bars indicate group means.

For transcript isoform composition (**Figure 2**), abundances are filtered and processed through IsoPlex to identify an effective number of isoforms [28, 29]. Genes with at least two effective isoforms proceed to feature construction (**Figure 2A**). Within each gene, isoform fractions are transformed into centered log-ratio coordinates, and smooth covariate-dependent trajectories are estimated. Singular value decomposition extracts the dominant fitted loading direction, which IsoVita approximates with the simplest near-optimal balance between two isoform groups, allowing other isoforms to remain unused (**Figure 2B**). The sample-level feature is the log ratio of the groups’ geometric mean fractions with interpretation determined by group membership.

**Figure 2:**
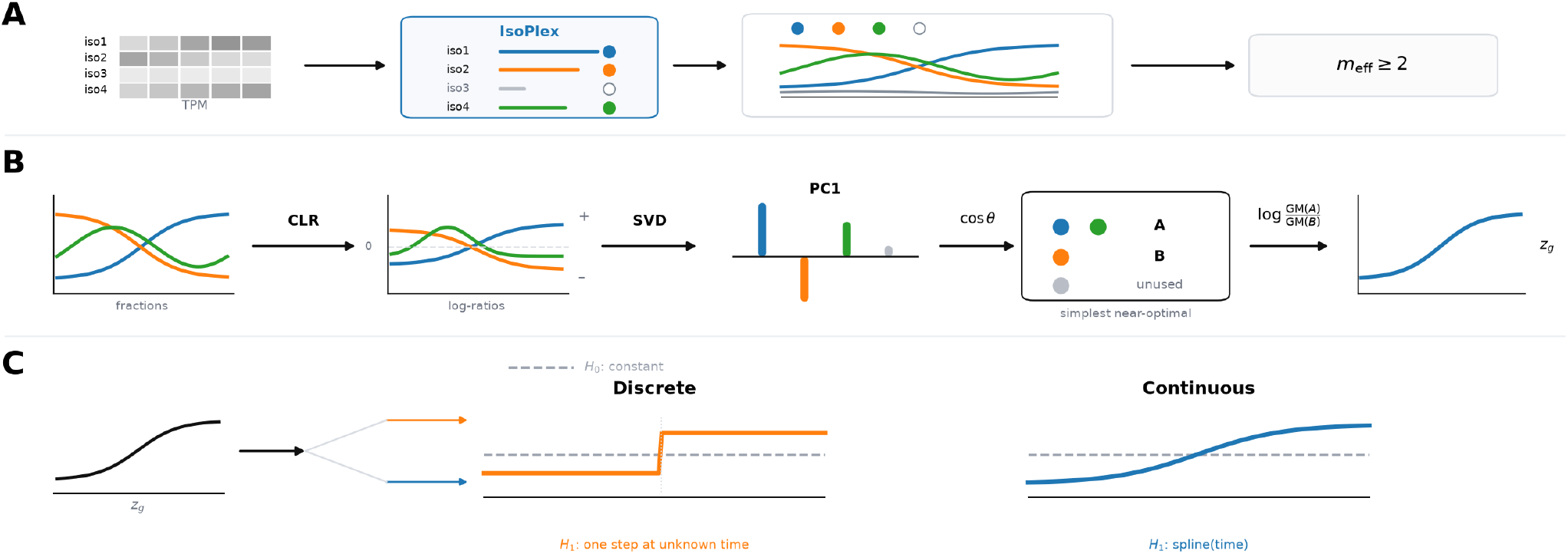
Overview of the IsoVita workflow. **(A)** Transcript abundances, expressed as transcripts per million (TPM), are processed through IsoPlex to identify effective isoforms; genes with at least two effective isoforms (*m*_eff_ ≥2) proceed to feature construction. **(B)** Within-gene isoform fractions undergo a centered log-ratio (CLR) transformation, followed by singular value decomposition (SVD) to identify the first principal component (PC1). Cosine similarity guides selection of a simple, near-optimal contrast between isoform groups A and B, with other isoforms left unused. The resulting gene-level feature is *z*_*g*_ = log[GM(*A*)*/*GM(*B*)], where GM denotes the geometric mean. **(C)** Two statistical inference branches test this feature against a constant null model: a discrete branch allowing one step at an unknown covariate value and a continuous branch modeling a B-spline function of the covariate.

Two branches test this feature against a constant null model: a discrete branch scans for a step at an unknown covariate value, and a continuous branch tests for a non-constant spline trajectory (**Figure 2C**). To account for covariate-informed feature selection, each permutation procedure repeats covariate-dependent feature construction and testing. Benjamini-Hochberg adjustment is applied separately to each branch where either or both branches may detect a gene. We next evaluated detection, false discoveries, and change point localization for the complete learning-and-testing procedure.

### 2.2 Temporal compositional variation with low empirical false discovery proportions

We first examined whether a compact, interpretable feature could retain the dominant pattern of temporal compositional variation. Across targeted simulations, the selected features closely approximated the dominant fitted direction, representing coordinated isoform changes through two opposing groups while allowing weakly aligned components to remain unused (**Figure S1**). These examples illustrate how feature condensation can simplify interpretation without requiring every isoform to contribute to the contrast. We then evaluated feature learning and inference together using transcript-level simulations containing temporally null genes and genes with step or sigmoid trajectories across a range of signal strengths (**Figure 3A, S2**). The mean empirical false discovery proportions remained below the nominal 5% level in both branches (**Figure 3B**), supporting the intended behavior of selection-aware inference despite using the covariate for both feature construction and testing. The union of branch discoveries also had a low empirical false discovery proportion, although it was not a separately FDR-controlled testing family.

**Figure 3:**
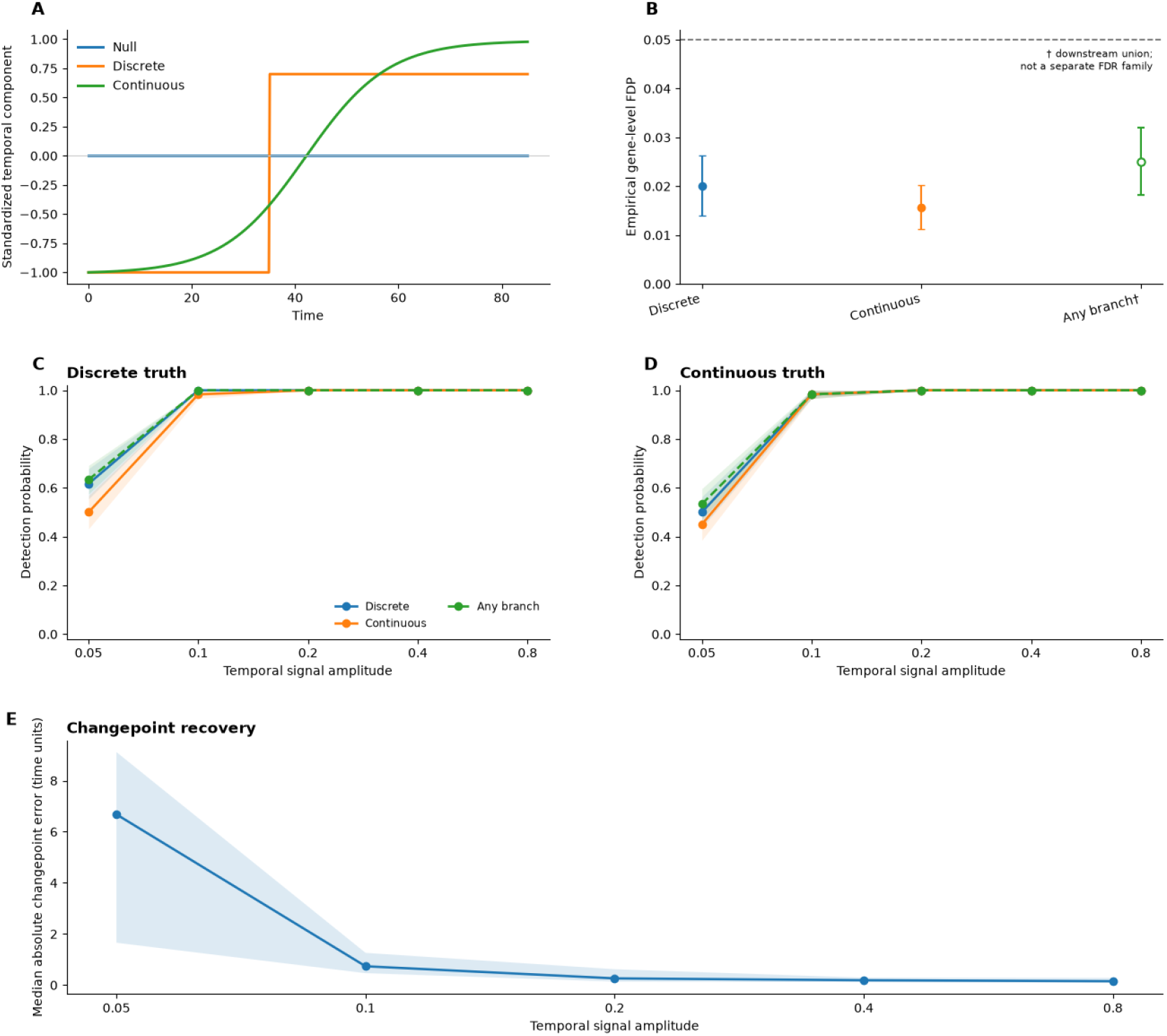
Evaluating IsoVita detection and change point recovery with synthetic data. **(A)** Standardized components for null, discrete, and continuous trajectories, with the discrete transition at time 35. **(B)** Mean empirical gene-level false discovery proportion (FDP) for each branch and their downstream union, with error bars showing ± 1 standard error across replicates. The dashed line marks the nominal 5% false discovery rate (FDR) threshold applied separately to each branch; the union is not a separately controlled FDR family. **(C)** Mean detection probability versus temporal signal amplitude for discrete truth; shading shows ±1 standard error across replicates. **(D)** Mean detection probability versus temporal signal amplitude for continuous truth; shading shows ±1 standard error across replicates. **(E)** Median across replicates of the within-replicate median absolute change point error, with shading indicating the interquartile range. Change point errors include all discrete-truth genes regardless of statistical significance.

Stronger temporal signals improved both the detection of compositional variation and the precision with which abrupt transitions could be located. Detection probability rose rapidly from partial recovery of weaker signals to nearly complete recovery as signal strength increased (**Figure 3C-D**). Increasing amplitude from 0.05 to 0.10 reduced median absolute change point error from approximately 6.7 to less than 1 time unit (**Figure 3E**), showing that stronger signals supported inference about both the presence and timing of a transition. Both branches frequently detected the same alternatives: a step can produce a non-constant spline fit, while a smooth transition can produce a substantial difference across a candidate split. Thus, the branches provide overlapping evidence of temporal association.

### 2.3 Trajectory shape and study design influence sensitivity to temporal variation

We next examined how performance depended on the form of the temporal signal. Weak steps near the boundaries of the sampled interval were harder to detect than those nearer its center, while stronger signals improved detection and change point localization across all tested locations (**Figure S3A-B**). The continuous branch detected both monotone and non-monotone trajectories, extending its sensitivity beyond the reference sigmoid pattern, although weak U-shaped signals were particularly challenging (**Figure S3C**). These differences diminished as signals strengthened. Together, these results show that IsoVita can detect a range of temporal patterns, but the signal strength needed for reliable detection depends on trajectory shape and transition location.

Study design and compositional complexity further influenced detection. Larger sample sizes and lower transcript-level noise improved power, while adding weakly informative isoforms around a fixed dominant contrast reduced sensitivity (**Figure S4A-C**). This decline occurred even though the learned feature could leave components unused, showing that this flexibility did not fully offset the loss of sensitivity associated with additional isoforms. Sampling coverage also mattered in addition to sample size (**Figure S4D**). Approximately balanced sampling yielded the highest detection rates in the settings examined, while early-heavy sampling retained sensitivity to the simulated step but substantially reduced detection of the later sigmoid transition. These results show that sensitivity depends on how sampling coverage aligns with the underlying temporal pattern. We next applied IsoVita to human brain RNA-sequencing data to examine age-associated isoform variation and the transcript contrasts underlying detected trajectories.

### 2.4 IsoVita identifies widespread and heterogeneous age-associated isoform trajectories in dlPFC

Previous studies identified widespread splicing changes during human brain development and aging, including shifts in *PALM* and *SNAP25* isoform usage in the dorsolateral prefrontal cortex (dlPFC) [25, 26]. We used this setting to examine whether IsoVita can connect established biological patterns to interpretable evidence of age association. Among 11,404 tested genes, 2,765 were detected by at least one branch at branch-specific FDR *<* 0.05 (**Figure 4A**), indicating that age-associated isoform variation extends beyond a few prominent examples.

**Figure 4:**
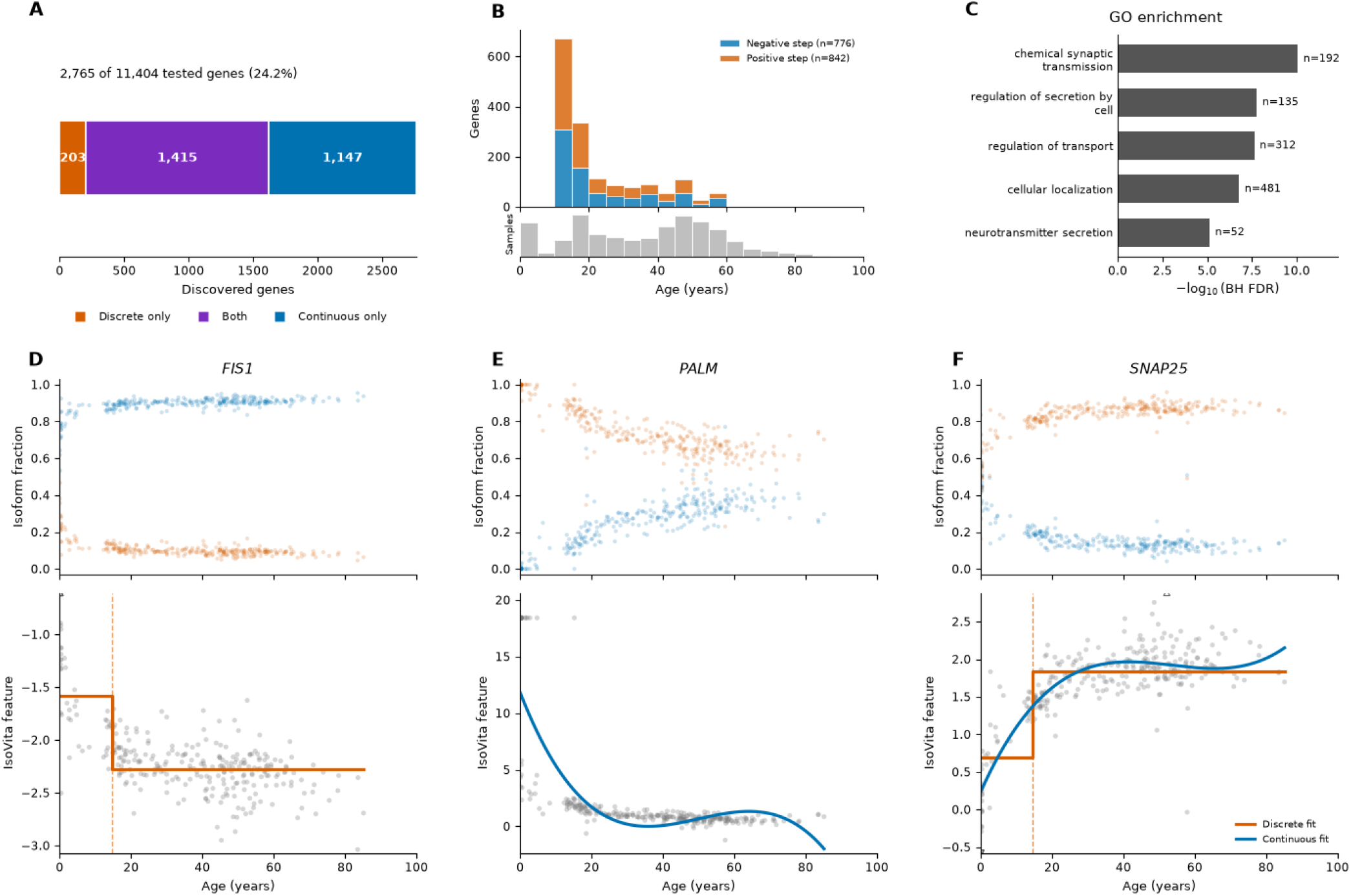
Age-associated isoform dynamics in the dorsolateral prefrontal cortex (dlPFC) dataset. **(A)** IsoVita identified 2,765 of 11,404 tested genes (24.2%): 203 were significant in the discrete branch only, 1,415 in both branches, and 1,147 in the continuous branch only. **(B)** Distribution of estimated change point ages among 1,618 discrete-branch discoveries, separated into negative (776 genes) and positive (842 genes) changes in the IsoVita feature; the lower histogram shows sample ages. **(C)** Five representative GO biological process terms enriched among the discovery union. Bars show ™ log_10_(BH FDR), and *n* indicates the number of uniquely mapped discovery genes annotated to each term. **(D-F)** Representative genes illustrating discrete-only (*FIS1*), continuous-only (*PALM*), and overlapping (*SNAP25*) discoveries. Upper plots show individual isoform fractions; lower plots show the condensed gene-level feature with discrete step fits in orange and continuous spline fits in blue. Dashed vertical lines mark estimated change points, and open triangles indicate observations beyond the displayed feature limits.

The substantial but incomplete overlap between discrete and continuous branches explains the value of testing more than one type of temporal pattern. Many genes were detected by both branches, as expected from the simulations: a significant age-associated change can support both a step approximation and a smooth trajectory. The overlap therefore reflects shared sensitivity to age association, rather than two distinct biological classes. However, 1,147 genes were detected only by the continuous branch, revealing that a single-step test would miss a substantial set of age associations.

Among genes detected by the discrete branch, estimated change points clustered in childhood and adolescence in dlPFC cohort (**Figure 4B**). Gene ontology (GO) enrichment further connected the discoveries to biological functions: enriched terms include chemical synaptic transmission, regulation of transport, and neurotransmitter secretion relative to the background containing all tested genes (**Figure 4C**).

*FIS1* was detected only by the discrete branch, with a fitted transition near age 15 (**Figure 4D**). Only the continuous branch detected the previously reported progressive shift in *PALM* isoform usage (**Figure 4E**). *SNAP25* was detected as significant by both branches, consistent with previous literature on significant developmental change in isoform usage (**Figure 4F**) [26]. These examples reveal how the branches capture overlapping evidence while preserving a direct connection between each association and contributing transcripts. IsoVita builds on earlier descriptions of isoform changes by providing a systematic way to learn the transcript contrasts and test their association with time without requiring a pre-specified single temporal pattern.

### 2.5 IsoVita extends interpretable isoform discovery to hippocampus

We next applied IsoVita to hippocampus data to examine whether the same workflow could support discovery in another brain region. Prior work established regional differences in expression and developmental regulation [27], motivating us to learn transcript contrast separately rather than reusing those identified in dlPFC. IsoVita identified 1,009 of 9,371 tested genes through at least one branch at branch-specific FDR *<* 0.05 (**Figure 5A**). Both branches discovered unique genes beyond their shared set, showing that the complementary value of the branches extended to a different regional context.

**Figure 5:**
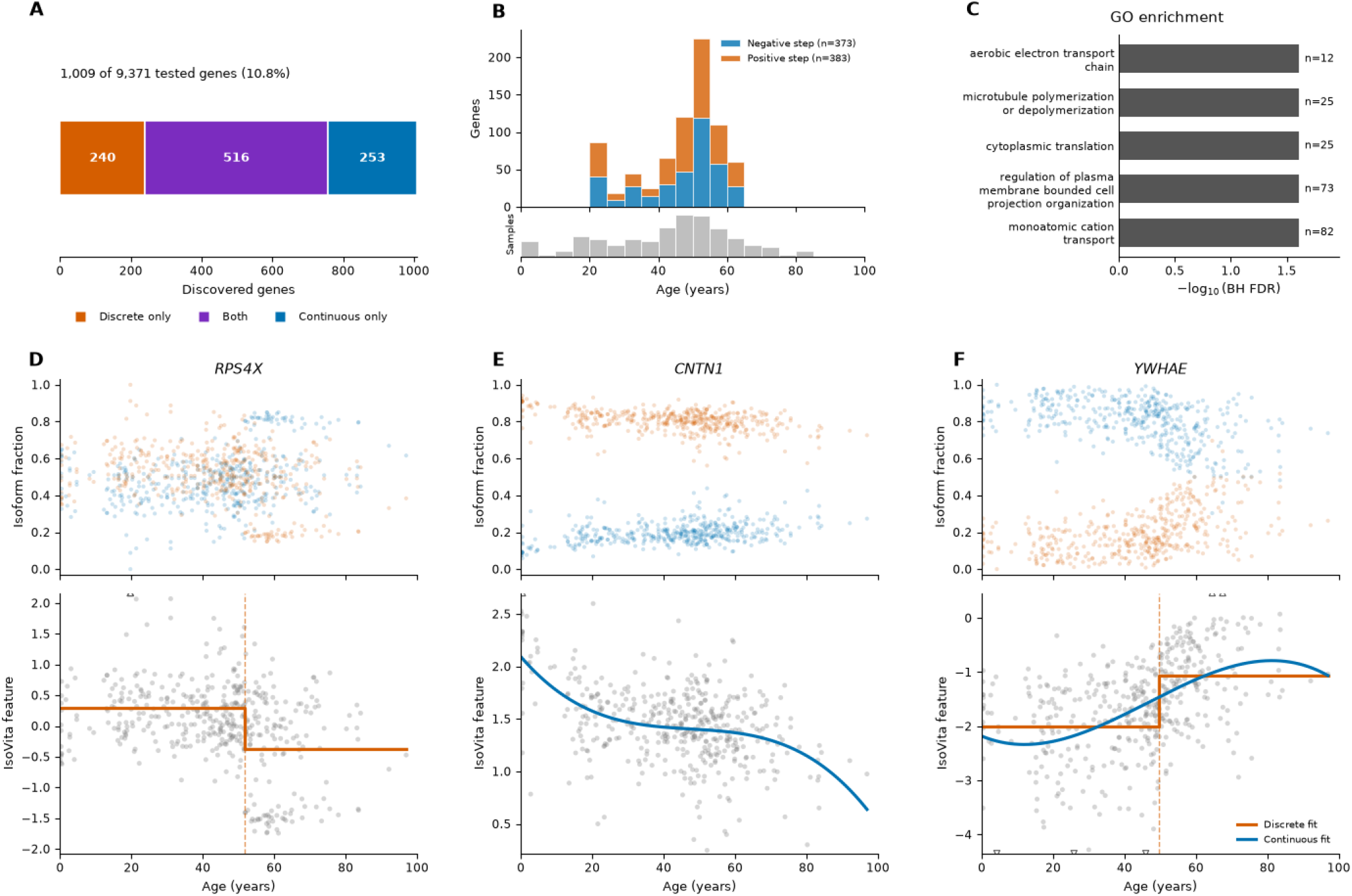
Age-associated isoform dynamics in the hippocampus dataset. **(A)** IsoVita identified 1,009 of 9,371 tested genes (10.8%): 240 were significant in the discrete branch only, 516 in both branches, and 253 in the continuous branch only. **(B)** Distribution of estimated change point ages among 756 discrete-branch discoveries, separated into negative (373 genes) and positive (383 genes) changes in the IsoVita feature; the lower histogram shows sample ages. **(C)** Five representative GO biological process terms enriched among the discovery union. Bars show ™ log_10_(BH FDR), and *n* indicates the number of uniquely mapped discovery genes annotated to each term. **(D-F)** Representative genes illustrating discrete-only (*RPS4X*), continuous-only (*CNTN1*), and overlapping (*YWHAE*) discoveries. Upper plots show individual isoform fractions; lower plots show the condensed gene-level feature with discrete step fits in orange and continuous spline fits in blue. Dashed vertical lines mark estimated change points, and open triangles indicate observations beyond the displayed feature limits.

Analysis in the hippocampus revealed a different temporal profile, with estimated change points concentrated near ages 50-55 rather than childhood and adolescence (**Figure 5B**). This contrast motivates further investigation of regional timing, although differences in sampling and detection sensitivity prevent a direct biological comparison. GO enrichment broadened the context beyond individual genes, identifying terms related to aerobic electron transport, microtubule dynamics, cytoplasmic translation, cell projection organization, and cation transport (**Figure 5C**). These enrichments suggest that age-associated isoform variation involves genes spanning energy metabolism, protein synthesis, and cellular organization, providing priorities for follow-up functional analyses.

*RPS4X* showed a discrete-branch association near age 52, *CNTN1* showed a continuous decline in its represented IsoVita feature, and *YWHAE* supported both step and smooth fits (**Figure 5D-F**). These learned contrasts turn gene-level discoveries into specific transcript groups that could be investigated further. For instance, earlier experiments linked *CNTN1* to hippocampal plasticity and memory in aged mice [30]. Here, the *CNTN1* result raises the additional question of how relative isoform usage contributes to its age-related biology.

While dlPFC connects IsoVita to previously established isoform variations, hippocampus demonstrates the use for generating interpretable candidates in another region. Together, the applications in individual brain regions support the use of a common learning-and-testing framework while allowing the contributing transcripts and temporal patterns to differ.

## 3 Discussion

IsoVita learns an interpretable compositional contrast and tests its association with a continuous covariate while accounting for feature learning. It combines a balance between two learned component groups, complementary change point and spline tests, and permutations that repeat feature construction and testing. This integration connects statistical evidence to explicit component membership. Although demonstrated using transcript isoforms, the statistical construction applies more broadly to compositions paired with a continuous covariate.

The simulations showed low empirical false discovery proportions within the settings examined. Detection power and change point accuracy improved with stronger temporal signals, while sensitivity depended on trajectory shape and sampling coverage. The brain RNA-sequencing applications connected gene-level discoveries to the contributing transcript groups, illustrating how IsoVita links evidence of age association to interpretable compositional contrasts. The two tests provide complementary evidence of covariate association, rather than definitive classifications of trajectory shape. An abrupt change may be detected by both tests, and a gradual trend may produce a significant change point result. The estimated change point is therefore a best-fitting step location, not proof of an abrupt underlying transition.

The current framework also has limits. One balance may miss additional modes of compositional variation, and sensitivity depends on how well the single-step and spline alternatives capture the underlying pattern. Repeating feature learning under permutation addresses selection, but does not remove confounding or measurement error. In bulk tissue, age-associated balances may reflect changes in cellular composition, changes in isoform usage within cell types, or both. Inference requires exchangeability under the null, so clustered, longitudinal, and covariate-adjusted designs need suitable modeling and resampling extensions. Applications beyond transcriptomics also require domain-appropriate preprocessing, particularly for uncertain measurements. Multiple balances, complex alternatives, and more efficient resampling are possible future directions for extending this approach to interpretable compositional inference.

## 4 Methods

### 4.1 Overview of IsoVita

IsoVita is a framework for detecting compositional variation along a continuous covariate while retaining an interpretable representation of the contributing components. For sample *i* = 1, …, *n*, gene *g*, and transcript *j*, let *x*_*igj*_ ≥ 0 denote transcript abundance and *t*_*i*_ the observed continuous covariate. The objective is to test whether within-gene isoform composition varies with *t*_*i*_ without requiring predefined transcript contrasts or changing the covariate into discrete classes.

The workflow consists of three stages (**Figure 2**). First, eligible isoforms are identified and condensed into a single gene-level log-ratio feature that approximates the dominant covariate-dependent compositional variation. Second, two complementary tests evaluate this feature for discrete change points and smooth continuous trajectories. Third, selection-aware permutations repeat feature learning and testing to account for covariate-guided feature construction. Multiple testing adjustment is performed separately for each branch, and their union is reported descriptively rather than as an independently FDR-controlled set.

### 4.2 Input preparation and effective isoform identification

For the transcript analyses, IsoVita takes a transcript-by-sample abundance table containing `gene_id, transcript_id`, and one finite, non-negative numeric abundance column per sample, together with matched sample metadata. Sample identifiers must be unique, and the abundance columns must match the metadata sample identifiers exactly and in the same order. Sample metadata contain `sample_id` and a numeric continuous covariate, named `age` in the preserved analysis code and `time` in the maintained package.

For the reported brain analyses, dataset-specific preparation used `prepare_isovita_inputs.py`. These steps removed samples with missing or non-numeric age, retained postnatal samples with age ≥ 0, preserved metadata order, merged transcript-to-gene annotations, removed unmapped transcripts, and verified finite, non-negative abundances. Postnatal restriction was specific to these applications. Input preparation does not convert raw counts to TPM; the supplied abundances must already be on the intended analysis scale.

For brain transcript analyses, effective isoform identification was performed after an expression eligibility filter: a transcript is retained if its abundance exceeds 0.5 TPM in at least five samples. After this transcript-level filter, genes are additionally required to have positive total abundance when summing over the retained transcripts, in every retained sample. IsoPlex is then applied to this filtered transcript table with expression type set to TPM to identify effective isoforms [28, 29]. *E*_*g*_ denotes the resulting effective-isoform set for gene *g* and *K*_*g*_ = |*E*_*g*_|: genes with *K*_*g*_ *<* 2 are excluded from downstream trajectory analysis. All genes with *K*_*g*_ ≥ 2 enter the same feature-construction procedure.

### 4.3 Construction of the IsoVita gene feature

#### 4.3.1 Compositional representation and covariate-dependent smoothing

For each eligible gene, only transcripts in *E*_*g*_ are retained for feature construction. Samples with a non-positive or non-finite effective isoform total are not usable for that gene, and at least 20 usable samples are required. For the gene-specific feature construction, testing, and permutation procedures below, usable samples are reindexed as *i* = 1, …, *n*, with *n* denoting the number of usable samples for the gene under consideration. For each usable sample, the abundance of each effective isoform is converted to a within-effective-set fraction:

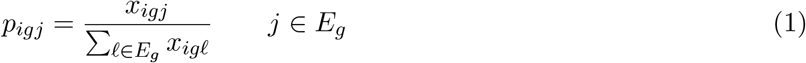

As logarithmic coordinates require strictly positive components, IsoVita adds a small pseudocount *E* = 10^™8^ to every component and then re-computes the composition as

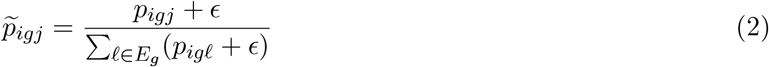

The stabilized composition is then mapped to centered log ratio (CLR) coordinates [3]. Specifically, the CLR representation removes the sample-specific compositional level and places the effective isoforms in a zero-sum log-ratio coordinate system.

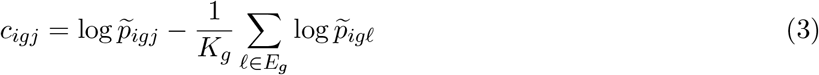

To learn the dominant covariate-dependent direction of variation, a cubic B-spline regression is fit separately to each CLR coordinate as a function of the covariate. Each regression includes an intercept and a cubic B-spline basis with four degrees of freedom, constructed using Patsy [31]. Let ĉ_*gj*_ (*t*) denote the fitted trajectory for effective isoform *j*. These trajectories are evaluated on a grid of *L* = 100 equally spaced covariate values spanning the observed covariate range for the gene. Each fitted trajectory is then centered over the evaluation grid, yielding the *L* × *K*_*g*_ matrix:

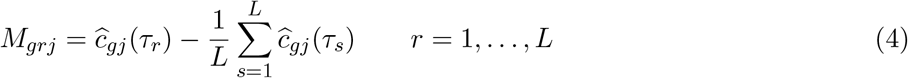

Centering across the grid removes fitted transcript-specific offsets so that subsequent decomposition is driven by temporal shape rather than baseline CLR values.

#### 4.3.2 Dominant temporal axis and group contrast approximation

IsoVita extracts the dominant fitted trajectory direction using the singular value decomposition (SVD) of the centered trajectory matrix:

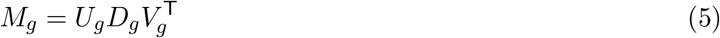

Let *v*_*g*_ denote the first column of *V*_*g*_, corresponding to the largest singular value. Its sign is fixed by requiring its largest-magnitude loading to be positive, and l*v*_*g*_l = 1.

The PC1 loading vector *v*_*g*_ is approximated by an interpretable two-group contrast. An assignment vector 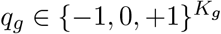 partitions effective isoforms into Group A (*q*_*gj*_ = +1), Group B (*q*_*gj*_ = ™1), and an optional unused set (*q*_*gj*_ = 0), with both groups non-empty. For group sizes *a* = |*A*_*g*_| and *b* = |*B*_*g*_|, the balance direction has weights proportional to 1*/a* for Group A, ™1*/b* for Group B, and zero otherwise. Let *d*(*q*_*g*_) denote the unit-norm direction [4, 17]. Candidate balances are evaluated by their absolute cosine similarity with PC1:

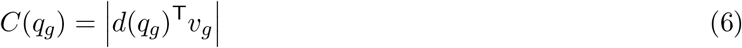

Rather than enumerating all 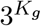 assignments, IsoVita uses an exact search over group sizes. For fixed (*a, b*), define the raw balance direction *w*_*A*,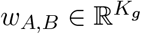_ as

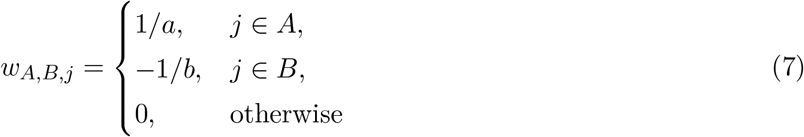

Its projection onto PC1 is

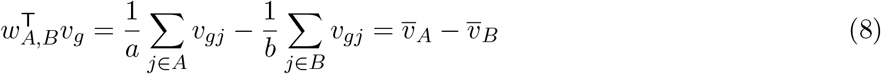

and its norm is

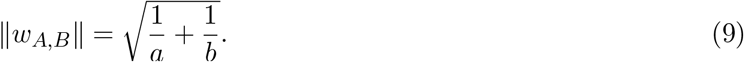

Since ‖*v*_*g*_ ‖ = 1, the signed cosine similarity is

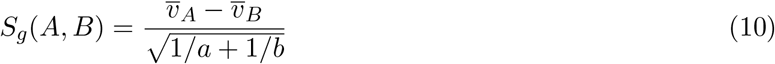

For fixed ordered group sizes (*a, b*), maximizing *S*_*g*_(*A, B*) is equivalent to maximizing 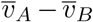. This difference is maximized by assigning the *a* largest entries of *v*_*g*_ to Group A and the *b* smallest entries to Group B: replacing any selected entry with a more extreme unselected entry can only increase the difference. Exchanging the two groups reverses the sign without changing the absolute cosine. Thus, IsoVita sorts *v*_*g*_ once and searches all total group sizes *m* = 2, …, *K*_*g*_ and ordered splits *a* = 1, …, *m*™1, with *b* = *m a*, to identify the maximum absolute cosine for each *m*.

Let 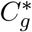 denote the globally maximum cosine similarity. With tolerance *δ* = 0.01, IsoVita selects the smallest total group size *m* whose best cosine is at least 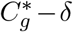, then chooses the highest-scoring assignment at that size. The final A/B orientation is chosen to have positive inner product with *v*_*g*_. For *K*_*g*_ = 2, the unique non-trivial balance is used directly. This procedure favors a simple, near-optimal approximation to the dominant fitted trajectory while allowing weakly aligned isoforms to remain unused.

#### 4.3.3 Definition and interpretation of the IsoVita feature

For the selected groups *A*_*g*_ and *B*_*g*_, the IsoVita feature for sample *i* and gene *g* is defined as:

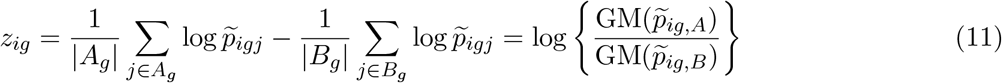

where GM denotes a geometric mean [16]. Therefore, positive and negative values indicate relative enrichment of Group A and Group B. The absolute direction is conventional because interchanging A and B multiplies the feature by ™1; F-statistics used in downstream inference are invariant to this sign change. Feature magnitude represents separation between the two learned isoform groups on a log-ratio scale and should not be interpreted as absolute gene expression.

### 4.4 Testing variation along the continuous covariate

After feature construction, each gene is represented by the vector *z*_*g*_ = (*z*_1*g*_, …, *z*_*ng*_)^T^. Two parallel tests compare a constant null model against either a single unknown step or a smooth spline trajectory. The resulting *F* -statistics are calibrated through the selection-aware permutation procedure described in Section 4.5, rather than through their classical *F* distributions.

#### 4.4.1 Discrete change point test

The discrete branch tests for a single step at an unknown covariate value. Samples are sorted by covariate, and a candidate split after position *s* is eligible only if adjacent covariate values differ and both sides contain at least

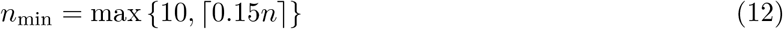

samples. Let RSS_0_ denote the residual sum of squares under the constant null model and RSS_1_(*s*) that under the step model. The split-specific statistic is

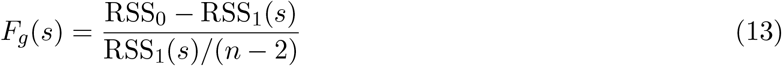

The observed discrete statistic maximizes over all eligible splits [32]:

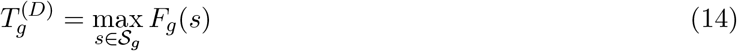

The estimated change point is the midpoint between the covariate values bracketing the maximizing split, with ties resolved by selecting the earliest split. The maximization is repeated under each permutation to account for change point selection.

#### 4.4.2 Continuous spline test

The continuous branch tests for a smooth, non-constant association with the covariate. The reduced model contains only an intercept, while the full model includes an intercept and a cubic B-spline basis with three degrees of freedom. Let *X*_0_ and *X*_1_ denote the reduced and full design matrices with ranks *r*_0_ and *r*_1_, and let RSS_0_ and RSS_1_ denote their residual sums of squares. The nested-model statistic is

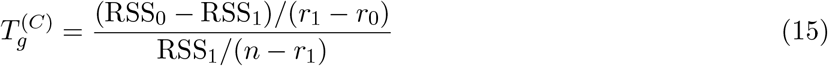

At least 20 samples are required. The spline alternative accommodates both monotone and non-monotone trajectories representable by the specified basis.

### 4.5 Selection-aware permutation inference and multiple testing

Feature learning in IsoVita depends on the covariate: the covariate determines the fitted spline trajectories, which in turn determine the dominant PC1 direction and selected group contrast. Because the contrast is learned using the same covariate subsequently tested for association, treating it as fixed would ignore this selection process and could lead to invalid inference [20]. IsoVita addresses this issue with a full pipeline covariate permutation procedure. Under the null of no association between the covariate and the gene’s isoform composition, covariate labels are permuted relative to the fixed composition data. For permutation *b*, the covariate-dependent feature learning procedure is re-executed, and the corresponding branch statistic is recomputed. Therefore, the permutation null includes both searching for a covariate-associated feature and evaluating the corresponding branch-specific test statistic [21].

Let *π*_*b*_ denote the *b*th permutation of the sample indices and let ℒ(*P*_*g*_, *t*) denote the complete feature-learning operation that maps the fixed effective-isoform composition *P*_*g*_ and a covariate vector *t* to an IsoVita feature. For genes with *K*_*g*_ *>* 2, the permuted feature is

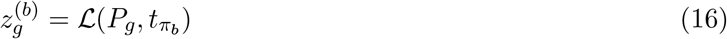

and the discrete or continuous statistic is recomputed using the same permuted covariate assignment. In practice, the composition and CLR coordinates are fixed, while every permutation refits the covariate-dependent spline trajectories, recomputes the trajectory SVD/PC1, relearns the group contrast, and reconstructs the feature before evaluating the requested detector. For *K*_*g*_ = 2, there is only one balance up to sign. Because F-statistics in both detection branches are invariant to feature sign, explicit re-learning is algebraically unnecessary in that special case; the implementation uses the equivalent reordered observed feature.

Let 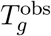 denote the observed detector statistic and 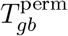 the full-pipeline statistic under permutation *b* = 1, …, *B*. The upper-tail Monte Carlo *p*-value uses the standard +1 correction,

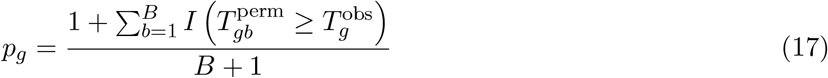

The minimum attainable value is therefore 1*/*(*B* + 1) [22]. The implementation issues a diagnostic warning when this resolution is coarser than the first nominal Benjamini–Hochberg (BH) threshold for a tested family.

BH adjustment is applied separately to all tested discrete hypotheses and all tested continuous hypotheses at nominal *α* = 0.05 [23]. No additional BH procedure is applied to the union of discoveries from two branches, and the union should therefore be interpreted as a descriptive combination of two branch-specific FDR analyses rather than as a separately calibrated third family.

### 4.6 Simulation framework

#### 4.6.1 Data generating model

Simulated data were generated at the transcript level before applying the IsoVita algorithm. This design avoids simulating data directly from the learned IsoVita feature. For simulated gene *g* with *K*_*g*_ transcripts, the latent log abundance of transcript *j* in sample *i* is

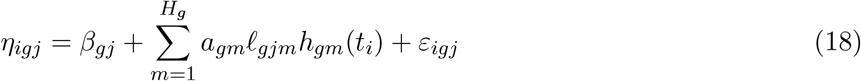

where *H*_*g*_ is the number of temporal components for gene *g, β*_*gj*_ is a transcript-specific baseline, *a*_*gm*_ is the amplitude of component *m*, _*gjm*_ is its loading for transcript *j*, and *h*_*gm*_(*t*) is its temporal basis function.

Each temporal basis is centered over the simulated time points and scaled to have maximum absolute value of 1. The simulator supports linear, sigmoid, step, U-shaped, Gaussian bump, and sinusoidal bases.

For a gene with *K*_*g*_ transcripts, the noise vector is Gaussian with covariance

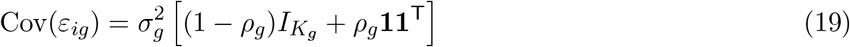

The latent transcript vector is converted to composition by a row-wise softmax function,

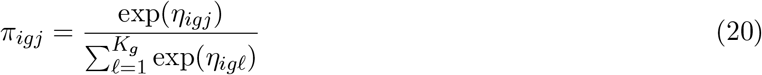

A positive gene-level abundance *G*_*ig*_ is drawn independently from a log-normal distribution with default log mean log(50) and log standard deviation 0.15, and transcript-level TPM-like values are generated as *x*_*igj*_ = *G*_*ig*_*π*_*igj*_. This gene-level scaling creates realistic variation in total gene abundance while preserving the simulated within-gene composition.

All simulated transcripts are marked effective in the simulated IsoPlex metrics. Therefore, the simulation studies mainly focus on stress-testing the IsoVita algorithm: they do not include evaluations of the IsoPlex effective-isoform selection algorithm.

#### 4.6.2 Primary simulation study

The primary simulation study was designed to evaluate IsoVita under the two trajectory patterns targeted by two different branches. Each simulation replicate contained 40 genes: 20 temporally null genes and 20 genes with temporal variation. The alternatives comprised two genes for every combination of two truth types (discrete and continuous) and five signal amplitudes (0.05, 0.10, 0.20, 0.40, and 0.80). Each gene contained four transcripts and each replicate contained *n* = 180 samples with time points drawn uniformly from 0 to 85 time units. Simulation time is expressed in arbitrary time units. 30 independent replicates were generated. Transcript baselines were sampled with standard deviation 0.22. Latent noise used parameters *σ* = 0.30 and *ρ* = 0.10.

Alternative genes were generated from a reference transcript loading direction (1.00, 0.82, −0.92, −0.75) with independent loading perturbations of standard deviation 0.05. The perturbed vector was centered and scaled before multiplication by the requested signal amplitude. Discrete alternatives used a step at time 35. Continuous alternatives used a sigmoid centered at time 42 with scale 16. Temporally null genes contained transcript-specific baselines and noise but no temporal component.

For each simulation replicate, IsoVita feature condensation was run using defaults, followed by parallel trajectory detectors. Each branch used *B* = 5000 permutations and nominal BH level 0.05. As all simulated transcripts are pre-specified as effective, IsoPlex is bypassed during all simulations.

Empirical false discovery proportion (FDP) was calculated separately for the discrete branch, continuous branch, and descriptive union. Within replicate *r* and detector *d*,

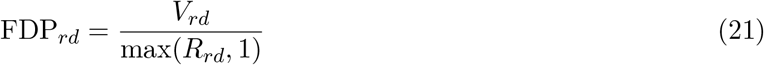

where *R*_*rd*_ is the number of discoveries and *V*_*rd*_ is the number of discovered genes generated under the complete temporally null model. As the two detection branches are overlapping detectors rather than trajectory classifiers, a continuous-truth gene detected by the discrete branch (or vice versa) is not counted as a false discovery: only temporally null genes are considered as false discoveries. Empirical power was calculated by truth type, signal amplitude, and detector as the proportion of corresponding non-null genes called significant within each replicate. Reported mean FDP and power curves average the replicate-level quantities. Pointwise 95% intervals were calculated as the replicate mean ± 1.96 standard errors and truncated to [0, 1].

change point localization was evaluated for every simulated discrete-truth gene, without conditioning on whether the gene was declared significant. For each gene, absolute localization error was 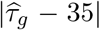. The median absolute error was first calculated within each replicate and signal-strength condition; these replicate-level medians were then summarized across replicates by their median and interquartile range. This evaluation separates change point estimation accuracy from the power threshold used to declare a discovery.

#### 4.6.3 Additional simulation studies

##### IsoVita feature construction stress tests

We constructed four targeted scenarios of increasing transcript-level complexity to evaluate whether the learned IsoVita feature resembled the dominant temporal variations. Scenarios ranges from a simple two-isoform contrast to multi-isoform settings containing secondary temporal structure. These experiments evaluated feature construction and were summarized by learned A/B/unused group assignments, number of retained isoforms, and cosine similarity to PC1.

##### Branch-specific trajectory geometry stress tests

We assessed the robustness of the discrete and continuous branches to alternative trajectory shapes beyond the primary simulation settings. Discrete alternatives varied the true change point location, while continuous alternatives included linear, sigmoid, U-shaped, and sinusoidal trajectories. Each setting was simulated across the same five signal amplitudes used in the primary study, with 15 replicates per condition. Only the corresponding detector was evaluated for each trajectory family. Inference used 1000 selection-aware permutations and BH correction at level 0.05. Performance was summarized by power (both branches) and change point localization error (discrete branch).

##### Background factor stress tests

We varied sample size, transcript-level noise, effective isoform count, and sampling-time distribution around a common reference simulation condition. The reference condition used signal amplitude 0.14, *n* = 80 samples, *K* = 4 effective isoforms, transcript-level noise standard deviation 0.35, *ρ* = 0.10, and uniformly sampled time points. In one-factor-at-a-time experiments, sample size varied over *n* ∈ {40, 60, 80, 120, 180}, noise standard deviation over *σ* ∈ {0.20, 0.28, 0.35, 0.45, 0.60}, and effective isoform count over *K* 2, 4, 6, 8, 12. Sampling time was varied among approximately balanced, uniform, middle-heavy, early-heavy, and late-heavy designs. Uniform time points were sampled from Unif(0, 85); middle-heavy, early-heavy, and late-heavy designs were generated by scaling Beta(2, 2), Beta(1.5, 2.5), and Beta(2.5, 1.5) draws to the interval [0, 85], while the balanced design used approximately equally spaced time points with small random perturbations. Each condition contained discrete, continuous, and null genes and was repeated 20 times. Both testing branches were run with 1000 selection-aware permutations and BH correction at level 0.05. Mean power and 95% confidence intervals, calculated as the mean ±1.96 standard errors across simulation replicates, were reported.

### 4.7 Real data analyses

IsoVita was applied independently to postnatal hippocampus and dorsolateral prefrontal cortex (dlPFC) bulk RNA-sequencing data. Dataset-specific processing was used only to obtain the canonical transcript-abundance and metadata inputs; all subsequent filtering, effective-isoform identification, feature construction, permutation inference, and branch-specific BH adjustment used the same IsoVita implementation and default statistical settings unless noted below.

#### 4.7.1 dlPFC

Bulk RNA-sequencing data and accompanying metadata for the dlPFC analysis were obtained from a previously published analysis of the dataset [26]. From the published analysis, we re-used transcript-level `dtu_scaled_tpm` abundance estimates generated using `tximport`, together with the corresponding sample metadata and transcript-to-gene annotation map [33]. Expression, metadata, and transcript-togene annotations were processed using `prepare_isovita_inputs_from_expression`. Preparation retained postnatal samples, removed unmapped transcripts, merged gene identifiers, and enforced exact sample alignment. The resulting dataset contained 285 samples, 157,310 mapped transcripts, and 57,921 genes spanning ages 0.01-85.14 years. Following effective isoform identification with IsoPlex, 11,404 genes with at least two effective isoforms entered IsoVita feature construction. dlPFC data were analyzed using both testing branches, 5000 selection-aware permutations, and branch-specific BH correction at level 0.05.

#### 4.7.2 Hippocampus

Bulk RNA-sequencing data and accompanying metadata for the hippocampus analysis were obtained from the LIBD BrainSeq Phase II resource [27]. Transcript models were taken from CHESS 3.1.3 after removal of alternative scaffolds and duplicate transcripts and used with the GRCh38 reference genome [34, 35]. Transcript sequences were extracted from the CHESS GTF using `gffread` [36]. A decoy-aware Salmon reference was constructed by concatenating transcriptome and genome sequences into a gentrome and using genomic sequence headers as decoys [37]. Salmon indexing used the –gencode option, and paired-end reads were quantified by selective alignment using –`validateMappings, –mimicBT2, –recoverOrphans`, and –`gcBias` [38]. A transcript-to-gene map was derived from the CHESS GTF, and sample-level Salmon outputs were imported with `pytximport`; the recorded downstream abundance representation was transcript-level `dtu_scaled_tpm` stored in condensed AnnData format [39, 40].

For the IsoVita analysis, the transcript-abundance AnnData object, sample metadata, and transcript-to-gene mapping were processed using `prepare_isovita_inputs`. Preparation retained hippocampus samples and postnatal ages, intersected expression and metadata, annotated transcripts with gene identifiers, and enforced exact sample ordering. The resulting dataset contained 442 samples, 157,458 mapped transcripts, and 57,960 genes spanning ages 0.07-96.92 years. Following effective isoform identification with IsoPlex, 9,371 genes with at least two effective isoforms entered IsoVita feature construction. Hippocampus data were analyzed using both testing branches, 5000 selection-aware permutations, and branch-specific BH correction at level 0.05.

#### 4.7.3 Gene ontology (GO) enrichment analysis

GO biological process enrichment was performed for each dataset using `g:Profiler` [41]. We tested genes detected by either IsoVita branch after branch-specific BH correction. The background included all genes that successfully completed IsoVita feature construction. CHESS identifiers were converted to Ensembl gene identifiers using GENCODE transcript annotations. Genes with no mapping or multiple mappings were excluded, and the enrichment background was restricted to mapped genes with GO biological process annotations. Terms with BH-adjusted p-values below 0.05 were considered significant. To reduce redundancy in the figures, we considered significant terms in order of increasing adjusted p-value. A term was retained only if its contributing discovery genes had a Jaccard similarity below 0.5 with those of every previously retained term. We displayed up to five retained terms.

### 4.8 Data and Software Availability

IsoVita is available as a public PyPI-installable package at https://pypi.org/project/isovita. The software source code is available on GitHub at https://github.com/danielchen05/isovita. The implementation used for all reported analyses is preserved independently of the maintained package at https://github.com/danielchen05/IsoVita-paper, together with preprocessing, analysis, and figure generation workflows. Processed bulk RNA-sequencing data inputs have been deposited in Zenodo at 10.5281/zenodo.23840334.

### 4.9 Acknowledgments, Funding, Authorship Contributions

## Acknowledgments

We thank the maintainers of the Joint High Performance Computing Exchange (JHPCE) compute cluster at Johns Hopkins Bloomberg School of Public Health for providing essential computing resources. We also thank Hicks lab for their constructive comments and suggestions. Further-more, we acknowledge the use of Artificial Intelligence to refine the language of the manuscript and assist with coding. The authors take full responsibility for its content and integrity.

## Author Contributions Statement

XC participated in project conceptualization, methodology, software, formal analysis, data curation, visualization, writing - original draft, and writing - review & editing. SCH participated in project conceptualization, supervision, writing - review & editing, and funding acquisition.

### Author ORCiD

## Supplementary Materials

### Contents

1. Supplementary Figures S1-S4

**Supplementary Figure S1:**
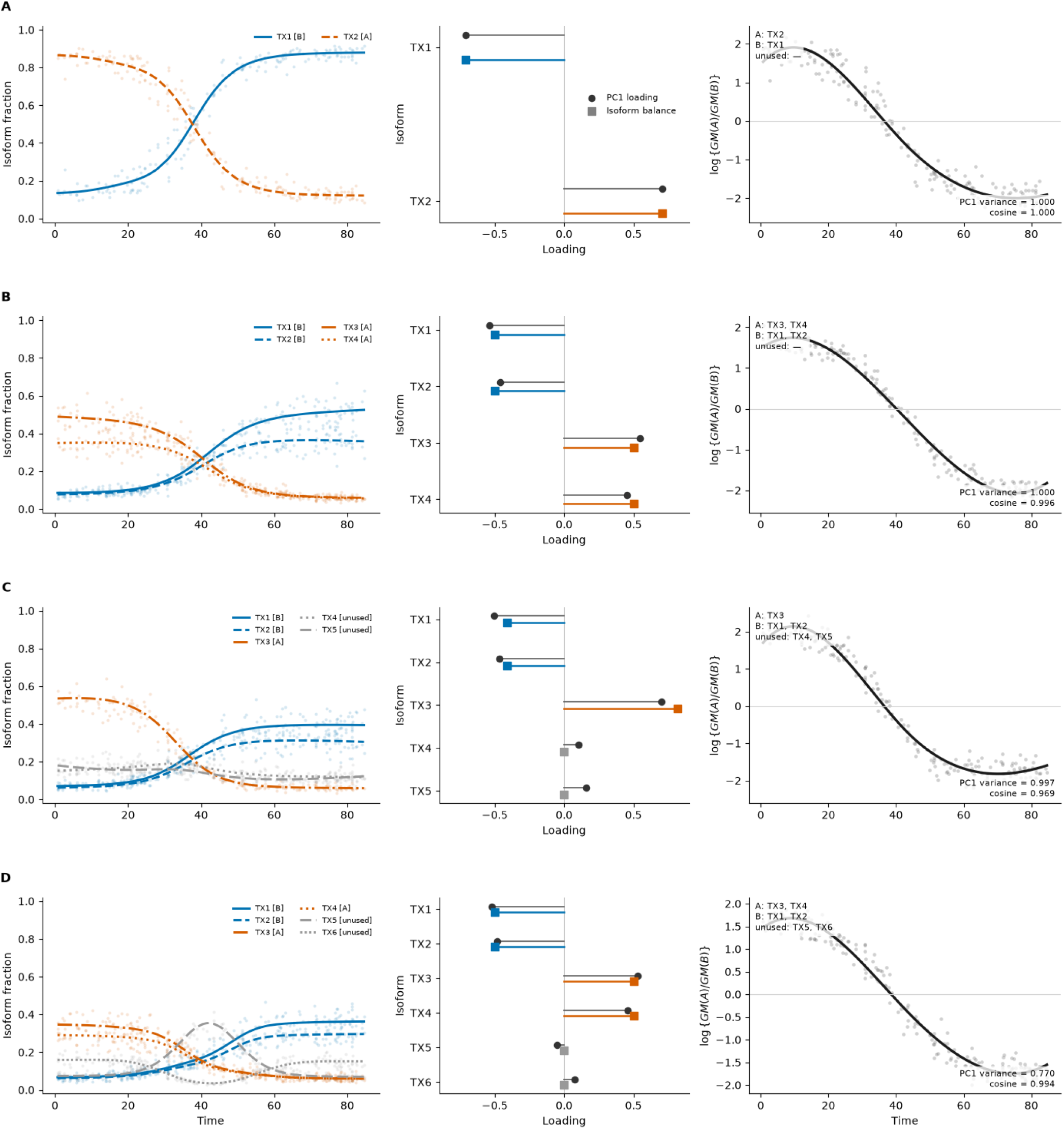
Condensation of simulated isoform trajectories into interpretable gene-level features. **(A)** A two-isoform contrast is represented exactly by the selected balance. **(B)** Four coherently changing isoforms are condensed into two opposing groups. **(C)** A dominant contrast between TX3 and TX1/TX2 is retained while TX4 and TX5 are excluded. **(D)** With competing temporal modes, the selected balance captures the dominant contrast between TX1/TX2 and TX3/TX4 while excluding TX5 and TX6. Annotations report the variance proportion explained by PC1 and the cosine similarity between PC1 and the selected direction represented by the two-group contrast. Each row shows observed isoform fractions and true underlying trajectories (left), PC1 loadings and the selected isoform-balance direction (middle), and the resulting log-ratio feature with a descriptive spline (right). Orange, blue, and gray denote group A, group B, and unused isoforms, respectively; circles and squares in the middle column denote PC1 and two-group contrast loadings.

**Supplementary Figure S2:**
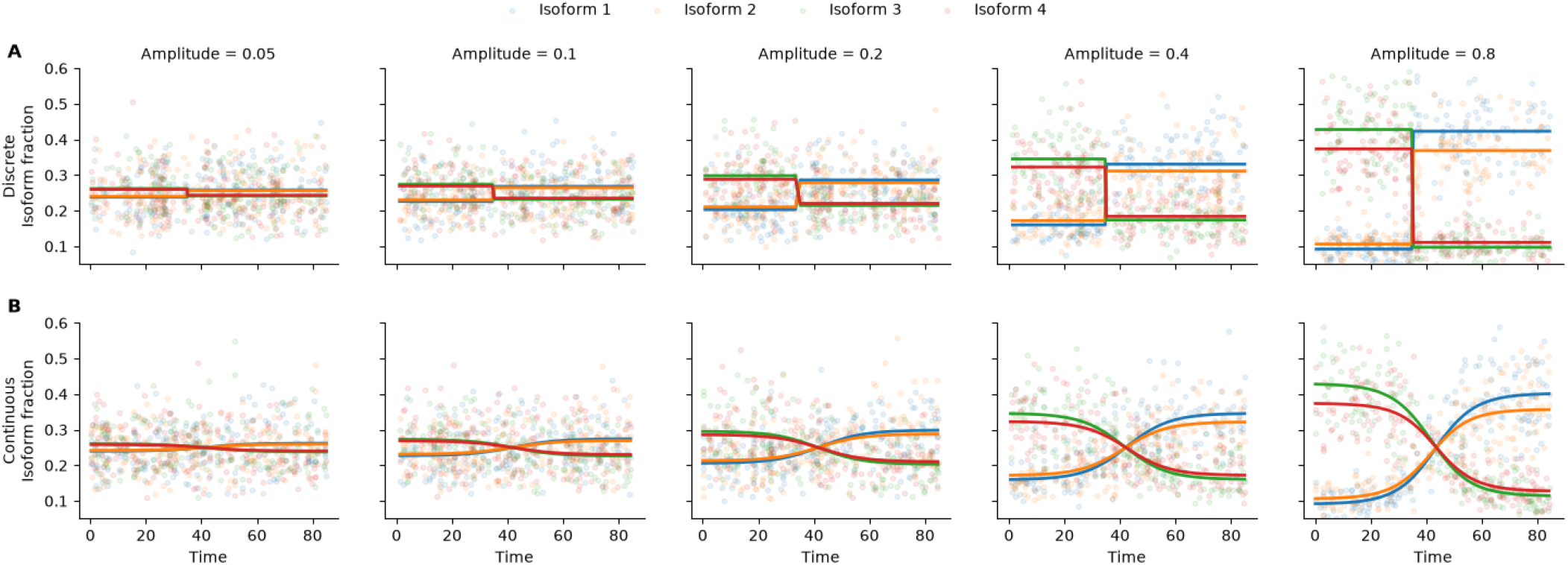
Simulated isoform fractions across temporal signal amplitudes. **(A)** Representative four-isoform genes with a discrete transition at time 35. **(B)** Representative four-isoform genes with continuous sigmoid trajectories. Columns show temporal signal amplitudes of 0.05, 0.10, 0.20, 0.40, and 0.80. Colored points represent noisy isoform fractions from 180 simulated samples per condition, and matching lines show the true underlying fraction trajectories. Increasing amplitude produces stronger reciprocal changes between isoforms under the same latent noise setting.

**Supplementary Figure S3:**
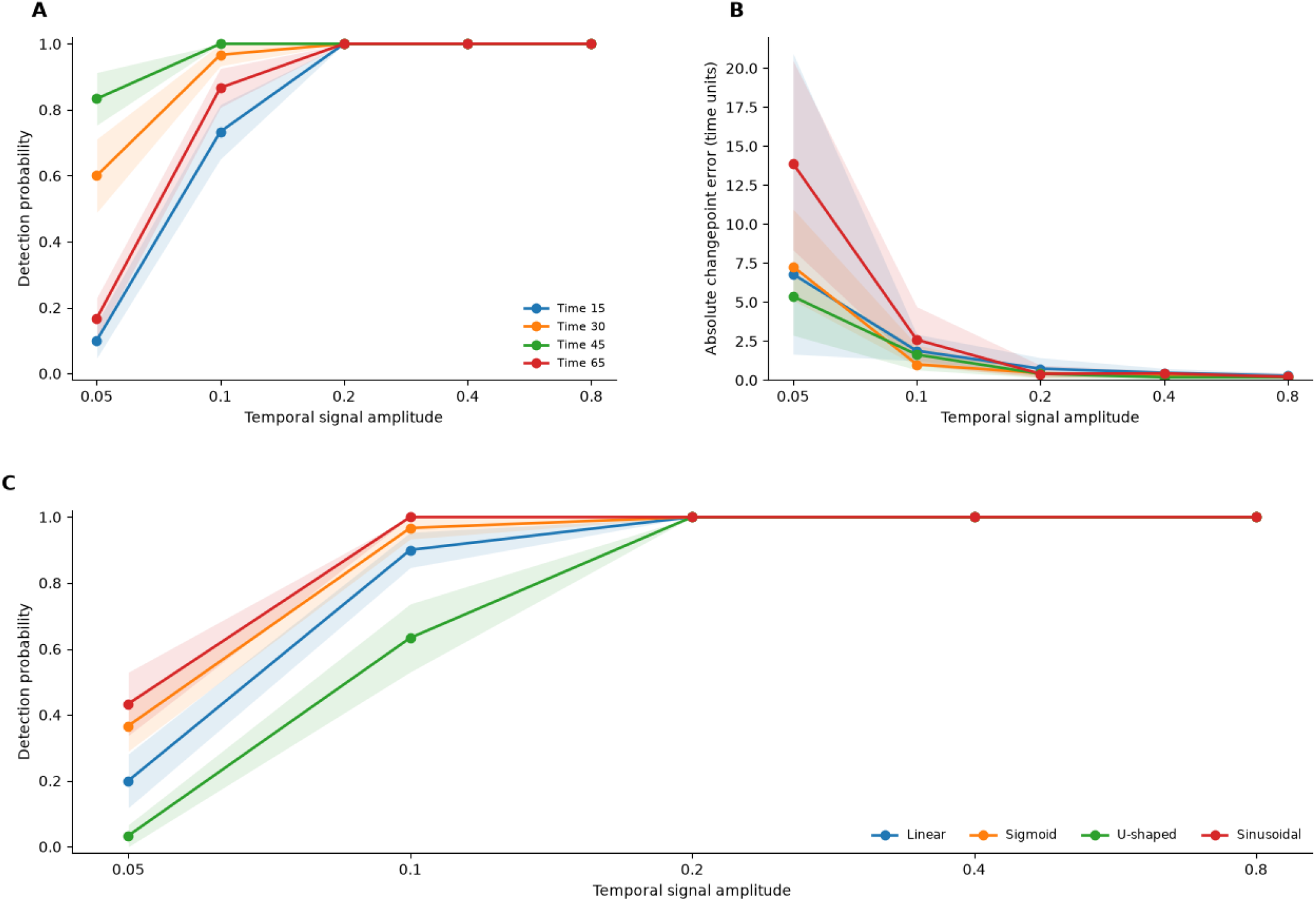
Sensitivity to change point location and continuous trajectory shape. Simulations used 15 replicates per temporal geometry, with 180 samples, four isoforms per gene, and equal numbers of null and alternative genes; discoveries were evaluated at a nominal 5% FDR within the corresponding branch. **(A)** Discrete-branch detection probability across signal amplitudes for true change points at time points 15, 30, 45, and 65. **(B)** Corresponding absolute change point errors, summarized by the median and interquartile range of replicate-level median errors across all discrete alternatives. **(C)** Continuous-branch detection probability for linear, sigmoid, U-shaped, and sinusoidal trajectories. Points and lines in A and C indicate mean detection probabilities, with shading showing ±1 standard error across replicates.

**Supplementary Figure S4:**
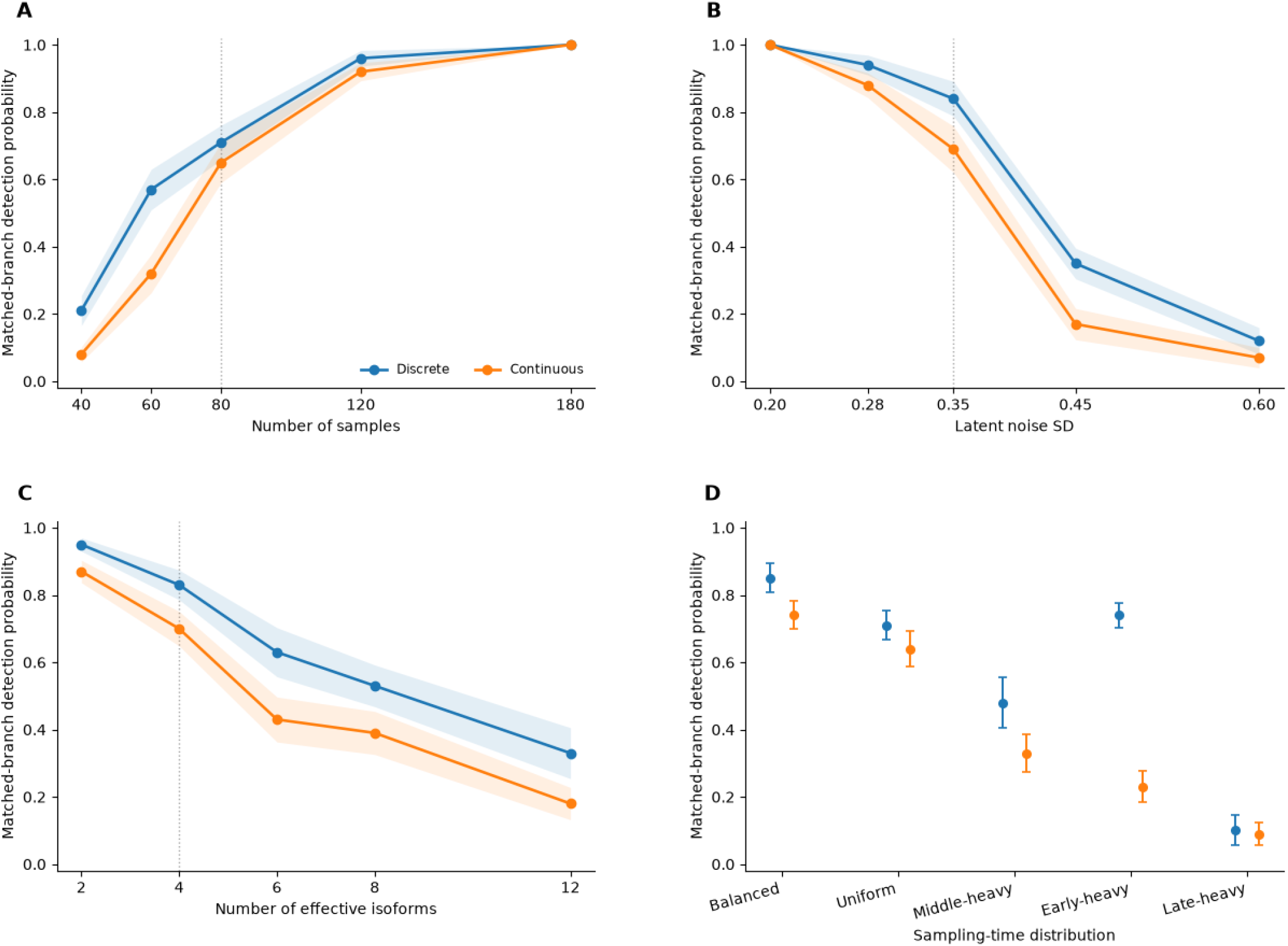
Sensitivity of IsoVita detection to sampling design, noise, and isoform complexity. Matched-branch detection probability denotes detection of discrete alternatives by the discrete branch and continuous alternatives by the continuous branch, each at a nominal 5% FDR. Simulations varied one factor at a time from a reference condition of 80 samples, four effective isoforms, latent noise standard deviation 0.35, uniformly distributed time points, and temporal signal amplitude 0.14. **(A)** Effect of sample size. **(B)** Effect of latent noise standard deviation. **(C)** Effect of increasing the number of effective isoforms while retaining two isoforms with strongest temporal signal and adding passenger isoforms. **(D)** Effect of balanced, uniform, middle-heavy, early-heavy, and late-heavy sampling-time distributions. Lines in (A)-(C) and points in (D) show mean detection probabilities across 20 replicates. Shading in (A)-(C) and error bars in (D) indicate ± 1 standard error. Sampling-time schemes are shown as separate categorical conditions without connecting lines. Dotted vertical lines in (A)-(C) mark reference settings.

## References

[1] K. Street, D. Risso, R. B. Fletcher, D. Das, J. Ngai, N. Yosef, E. Purdom, and S. Dudoit. Slingshot: cell lineage and pseudotime inference for single-cell transcriptomics. BMC Genomics, 19(1):477, Dec. 2018. ISSN 1471-2164. doi:10.1186/s12864-018-4772-0. URL https://bmcgenomics.biomedcentral.com/articles/10.1186/s12864-018-4772-0.

[2] K. Van Den Berge, H. Roux D. Bézieux, K. Street, W. Saelens, R. Cannoodt, Y. Saeys, S. Dudoit, and L. Clement. Trajectory-based differential expression analysis for single-cell sequencing data. Nature Communications, 11(1):1201, Mar. 2020. ISSN 2041-1723. doi:10.1038/s41467-020-14766-3. URL https://www.nature.com/articles/s41467-020-14766-3.

[3] J. Aitchison. The Statistical Analysis of Compositional Data. Journal of the Royal Statistical Society Series B: Statistical Methodology, 44(2):139–160, Jan. 1982. ISSN 1369-7412, 1467-9868. doi:10.1111/j.2517-6161.1982.tb01195.x. URL https://academic.oup.com/jrsssb/article/44/2/139/7027742.

[4] J. J. Egozcue, V. Pawlowsky-Glahn, G. Mateu-Figueras, and C. Barceló-Vidal. Isometric Logratio Transformations for Compositional Data Analysis. Mathematical Geology, 35(3):279–300, Apr. 2003. ISSN 1573-8868. doi:10.1023/A:1023818214614. URL https://doi.org/10.1023/A:1023818214614.

[5] E. T. Wang, R. Sandberg, S. Luo, I. Khrebtukova, L. Zhang, C. Mayr, S. F. Kingsmore, G. P. Schroth, and C. B. Burge. Alternative isoform regulation in human tissue transcriptomes. Nature, 456(7221):470–476, Nov. 2008. ISSN 0028-0836, 1476-4687. doi:10.1038/nature07509. URL https://www.nature.com/articles/nature07509.

[6] S. Anders, A. Reyes, and W. Huber. Detecting differential usage of exons from RNA-seq data. Genome Research, 22(10):2008–2017, Oct. 2012. ISSN 1088-9051. doi:10.1101/gr.133744.111. URL https://genome.cshlp.org/content/22/10/2008.

[7] M. Nowicka and M. D. Robinson. DRIMSeq: a Dirichlet-multinomial framework for multivariate count outcomes in genomics. F1000Research, 5:1356, Dec. 2016. ISSN 2046-1402. doi:10.12688/f1000re-search.8900.2. URL https://f1000research.com/articles/5-1356/v2.

[8] Y. I. Li, D. A. Knowles, J. Humphrey, A. N. Barbeira, S. P. Dickinson, H. K. Im, and J. K. Pritchard. Annotation-free quantification of RNA splicing using LeafCutter. Nature Genetics, 50(1):151–158, Jan. 2018. ISSN 1546-1718. doi:10.1038/s41588-017-0004-9. URL https://www.nature.com/articles/s41588-017-0004-9.

[9] B. Erdogdu, A. Varabyou, S. C. Hicks, S. L. Salzberg, and M. Pertea. Detecting differential transcript usage in complex diseases with SPIT. Cell Reports Methods, 4(3):100736, Mar. 2024. ISSN 26672375. doi:10.1016/j.crmeth.2024.100736. URL https://linkinghub.elsevier.com/retrieve/pii/S2667237524000614.

[10] W. Sun, Y. Liu, J. J. Crowley, T.-H. Chen, H. Zhou, H. Chu, S. Huang, P.-F. Kuan, Y. Li, D. Miller, G. Shaw, Y. Wu, V. Zhabotynsky, L. McMillan, F. Zou, P. F. Sullivan, and F. P.-M. De Villena. IsoDOT Detects Differential RNA-Isoform Expression/Usage With Respect to a Categorical or Continuous Covariate With High Sensitivity and Specificity. Journal of the American Statistical Association, 110(511):975–986, July 2015. ISSN 0162-1459, 1537-274X. doi:10.1080/01621459.2015.1040880. URL http://www.tandfonline.com/doi/full/10.1080/01621459.2015.1040880.

[11] A. M. Young, S. Van Buren, and N. U. Rashid. Differential transcript usage analysis incorporating quantification uncertainty via compositional measurement error regression modeling. Biostatistics, 25 (2):559–576, Apr. 2024. ISSN 1465-4644, 1468-4357. doi:10.1093/biostatistics/kxad008. URL https://academic.oup.com/biostatistics/article/25/2/559/7115316.

[12] J. Gilis, K. Vitting-Seerup, K. Van Den Berge, and L. Clement. satuRn: Scalable analysis of differential transcript usage for bulk and single-cell RNA-sequencing applications. F1000Research, 10:374, Aug. 2022. ISSN 2046-1402. doi:10.12688/f1000research.51749.2. URL https://f1000research.com/articles/10-374/v2.

[13] H. Topa and A. Honkela. Analysis of differential splicing suggests different modes of short-term splicing regulation. Bioinformatics, 32(12):i147–i155, June 2016. ISSN 1367-4811, 1367-4803. doi:10.1093/bioinformatics/btw283. URL https://academic.oup.com/bioinformatics/article/32/12/i147/2289066.

[14] M. J. Nueda, J. Martorell-Marugan, C. Martí, S. Tarazona, and A. Conesa. Identification and visualization of differential isoform expression in RNA-seq time series. Bioinformatics, 34(3):524–526, Feb. 2018. ISSN 1367-4803, 1367-4811. doi:10.1093/bioinformatics/btx578. URL https://academic.oup.com/bioinformatics/article/34/3/524/4158165.

[15] A. Bennett, E. K. Johnson, and K. J. M. Benjamin. IsoGraph resolves coordinated transcript choice and gene abundance in the aging human brain, Oct. 2026. URL http://biorxiv.org/lookup/doi/10.64898/2026.10.04.756439.

[16] J. Rivera-Pinto, J. J. Egozcue, V. Pawlowsky-Glahn, R. Paredes, M. Noguera-Julian, and M. L. Calle. Balances: a New Perspective for Microbiome Analysis. mSystems, 3(4):10.1128/msystems.00053–18, Aug. 2018. ISSN 2379-5077. doi:10.1128/msystems.00053-18. URL https://journals.asm.org/doi/10.1128/msystems.00053-18.

[17] J. A. Martín-Fernández, V. Pawlowsky-Glahn, J. J. Egozcue, and R. Tolosona-Delgado. Advances in Principal Balances for Compositional Data. Mathematical Geosciences, 50(3):273–298, Apr. 2018. ISSN 1874-8953. doi:10.1007/s11004-017-9712-z. URL https://doi.org/10.1007/s11004-017-9712-z.

[18] M. L. Calle, M. Pujolassos, and A. Susin. coda4microbiome: compositional data analysis for microbiome cross-sectional and longitudinal studies. BMC Bioinformatics, 24(1):82, Mar. 2023. ISSN 1471-2105. doi:10.1186/s12859-023-05205-3. URL https://bmcbioinformatics.biomedcentral.com/articles/10.1186/s12859-023-05205-3.

[19] R. Lu, G. Yu, C. Zhang, and Z. Chen. Sparse time-varying log-ratios for longitudinal high-throughput sequencing data. Frontiers in Bioinformatics, 6:1824265, June 2026. ISSN 2673-7647. doi:10.3389/fbinf.2026.1824265. URL https://www.frontiersin.org/articles/10.3389/fbinf.2026.1824265/full.

[20] N. Kriegeskorte, W. K. Simmons, P. S. F. Bellgowan, and C. I. Baker. Circular analysis in systems neuroscience: the dangers of double dipping. Nature Neuroscience, 12(5):535–540, May 2009. ISSN 1097-6256, 1546-1726. doi:10.1038/nn.2303. URL https://www.nature.com/articles/nn.2303.

[21] J. Hemerik and J. Goeman. Exact testing with random permutations. TEST, 27(4):811–825, Dec. 2018. ISSN 1133-0686, 1863-8260. doi:10.1007/s11749-017-0571-1. URL http://link.springer.com/10.1007/s11749-017-0571-1.

[22] B. Phipson and G. K. Smyth. Permutation P-values Should Never Be Zero: Calculating Exact P-values When Permutations Are Randomly Drawn. Statistical Applications in Genetics and Molecular Biology, 9(1), Oct. 2010. ISSN 1544-6115. doi:10.2202/1544-6115.1585. URL https://www.degruyterbrill.com/document/doi/10.2202/1544-6115.1585/html.

[23] Y. Benjamini and Y. Hochberg. Controlling the False Discovery Rate: A Practical and Powerful Approach to Multiple Testing. Journal of the Royal Statistical Society Series B: Statistical Methodology, 57(1):289–300, Jan. 1995. ISSN 1369-7412, 1467-9868. doi:10.1111/j.2517-6161.1995.tb02031.x. URL https://academic.oup.com/jrsssb/article/57/1/289/7035855.

[24] H. J. Kang, Y. I. Kawasawa, F. Cheng, Y. Zhu, X. Xu, M. Li, A. M. M. Sousa, M. Pletikos, K. A. Meyer, G. Sedmak, T. Guennel, Y. Shin, M. B. Johnson, Krsnik, S. Mayer, S. Fertuzinhos, S. Umlauf, S. N. Lisgo, A. Vortmeyer, D. R. Weinberger, S. Mane, T. M. Hyde, A. Huttner, M. Reimers, J. E. Kleinman, and N. estan. Spatio-temporal transcriptome of the human brain. Nature, 478(7370): 483–489, Oct. 2011. ISSN 1476-4687. doi:10.1038/nature10523. URL https://www.nature.com/articles/nature10523.

[25] P. Mazin, J. Xiong, X. Liu, Z. Yan, X. Zhang, M. Li, L. He, M. Somel, Y. Yuan, Y. Phoebe Chen, N. Li, Y. Hu, N. Fu, Z. Ning, R. Zeng, H. Yang, W. Chen, M. Gelfand, and P. Khaitovich. Widespread splicing changes in human brain development and aging. Molecular Systems Biology, 9(1):MSB201267, Jan. 2013. ISSN 1744-4292. doi:10.1038/msb.2012.67. URL https://doi.org/10.1038/msb.2012.67.

[26] B. Erdogdu, H. J. Ji, Z. C. Rudnick, M. Pertea, and S. L. Salzberg. Isoswitching in human brain development and aging, May 2025. URL http://biorxiv.org/lookup/doi/10.1101/2025.05.05.652255.

[27] L. Collado-Torres, E. E. Burke, A. Peterson, J. Shin, R. E. Straub, A. Rajpurohit, S. A. Semick, W. S. Ulrich, A. J. Price, C. Valencia, R. Tao, A. Deep-Soboslay, T. M. Hyde, J. E. Kleinman, D. R. Weinberger, and A. E. Jaffe. Regional Heterogeneity in Gene Expression, Regulation, and Coherence in the Frontal Cortex and Hippocampus across Development and Schizophrenia. Neuron, 103(2):203–216.e8, July 2019. ISSN 08966273. doi:10.1016/j.neuron.2019.05.013. URL https://linkinghub.elsevier.com/retrieve/pii/S0896627319304386.

[28] M. D. Schertzer, S. H. Park, J. Su, F. Reese, G. M. Sheynkman, and D. A. Knowles. Perplexity as a metric for isoform diversity in the human transcriptome. Genome Biology, Sept. 2026. ISSN 1474-760X. doi:10.1186/s13059-026-04260-2. URL https://link.springer.com/10.1186/s13059-026-04260-2.

[29] Fairlie Reese. fairliereese/isoplex: 0.1, Dec. 2025. URL https://zenodo.org/doi/10.5281/zenodo.18035396.

[30] D. Puzzo, A. Bizzoca, C. Loreto, C. A. Guida, W. Gulisano, G. Frasca, M. Bellomo, S. Castorina, G. Gennarini, and A. Palmeri. Role of F3/contactin expression profile in synaptic plasticity and memory in aged mice. Neurobiology of Aging, 36(4):1702–1715, Apr. 2015. ISSN 01974580. doi:10.1016/j.neurobiolaging.2015.01.004. URL https://linkinghub.elsevier.com/retrieve/pii/S019745801500007X.

[31] Nathaniel J. Smith, Matthew Wardrop, Christian Hudon, broessli, Peter Quackenbush, Skipper Seabold, Andrey Portnoy, Ben Beasley, Cameron Davidson-Pilon, Hassan Kibirige, Katrin Leinweber, Kevin Sheppard, Max Humber, Maximilien Colange, Michael Hudson-Doyle, Milo Koreniak, and Tim Gates. pydata/patsy: v0.5.6, Jan. 2024. URL https://zenodo.org/doi/10.5281/zenodo.592075.

[32] D. W. K. Andrews. Tests for Parameter Instability and Structural Change With Unknown Change Point. Econometrica, 61(4):821, July 1993. ISSN 00129682. doi:10.2307/2951764. URL https://www.jstor.org/stable/2951764?origin=crossref.

[33] C. Soneson, M. I. Love, and M. D. Robinson. Differential analyses for RNA-seq: transcript-level estimates improve gene-level inferences. F1000Research, 4:1521, Feb. 2016. ISSN 2046-1402. doi:10.12688/f1000research.7563.2. URL https://f1000research.com/articles/4-1521/v2.

[34] A. Varabyou, M. J. Sommer, B. Erdogdu, I. Shinder, I. Minkin, K.-H. Chao, S. Park, J. Heinz, C. Pockrandt, A. Shumate, N. Rincon, D. Puiu, M. Steinegger, S. L. Salzberg, and M. Pertea. CHESS 3: an improved, comprehensive catalog of human genes and transcripts based on large-scale expression data, phylogenetic analysis, and protein structure. Genome Biology, 24(1):249, Oct. 2023. ISSN 1474-760X. doi:10.1186/s13059-023-03088-4. URL https://genomebiology.biomedcentral.com/articles/10.1186/s13059-023-03088-4.

[35] V. A. Schneider, T. Graves-Lindsay, K. Howe, N. Bouk, H.-C. Chen, P. A. Kitts, T. D. Murphy, K. D. Pruitt, F. Thibaud-Nissen, D. Albracht, R. S. Fulton, M. Kremitzki, V. Magrini, C. Markovic, S. McGrath, K. M. Steinberg, K. Auger, W. Chow, J. Collins, G. Harden, T. Hubbard, S. Pelan, J. T. Simpson, G. Threadgold, J. Torrance, J. M. Wood, L. Clarke, S. Koren, M. Boitano, P. Peluso, H. Li, C.-S. Chin, A. M. Phillippy, R. Durbin, R. K. Wilson, P. Flicek, E. E. Eichler, and D. M. Church. Evaluation of GRCh38 and de novo haploid genome assemblies demonstrates the enduring quality of the reference assembly. Genome Research, 27(5):849–864, May 2017. ISSN 1088-9051, 1549-5469. doi:10.1101/gr.213611.116. URL http://genome.cshlp.org/lookup/doi/10.1101/gr.213611.116.

[36] G. Pertea and M. Pertea. GFF Utilities: GffRead and GffCompare. F1000Research, 9:304, Sept. 2020. ISSN 2046-1402. doi:10.12688/f1000research.23297.2. URL https://f1000research.com/articles/9-304/v2.

[37] R. Patro, G. Duggal, M. I. Love, R. A. Irizarry, and C. Kingsford. Salmon provides fast and bias-aware quantification of transcript expression. Nature Methods, 14(4):417–419, Apr. 2017. ISSN 1548-7091, 1548-7105. doi:10.1038/nmeth.4197. URL https://www.nature.com/articles/nmeth.4197.

[38] A. Srivastava, L. Malik, H. Sarkar, M. Zakeri, F. Almodaresi, C. Soneson, M. I. Love, C. Kingsford, and R. Patro. Alignment and mapping methodology influence transcript abundance estimation. Genome Biology, 21(1):239, Dec. 2020. ISSN 1474-760X. doi:10.1186/s13059-020-02151-8. URL https://genomebiology.biomedcentral.com/articles/10.1186/s13059-020-02151-8.

[39] M. Kuehl, M. N. Wong, N. Wanner, S. Bonn, and V. G. Puelles. Gene count estimation with pytximport enables reproducible analysis of bulk RNA sequencing data in Python. Bioinformatics, 40(12): btae700, Nov. 2024. ISSN 1367-4811. doi:10.1093/bioinformatics/btae700. URL https://academic.oup.com/bioinformatics/article/doi/10.1093/bioinformatics/btae700/7905459.

[40] I. Virshup, S. Rybakov, F. J. Theis, P. Angerer, and F. A. Wolf. anndata: Access and store annotated datamatrices. Journal of Open Source Software, 9(101):4371, Sept. 2024. ISSN 2475-9066. doi:10.21105/joss.04371. URL https://joss.theoj.org/papers/10.21105/joss.04371.

[41] L. Kolberg, U. Raudvere, I. Kuzmin, P. Adler, J. Vilo, and H. Peterson. g:Profilerinteroperable web service for functional enrichment analysis and gene identifier mapping (2023 update). Nucleic Acids Research, 51(W1):W207–W212, July 2023. ISSN 0305-1048, 1362-4962. doi:10.1093/nar/gkad347. URL https://academic.oup.com/nar/article/51/W1/W207/7152869.

